# Activity based proteome profiling of serum serine hydrolases: application in pediatric abusive head trauma

**DOI:** 10.1101/2024.04.17.589869

**Authors:** Estelle Maret, Kim Wiskott, Tobias Shipley, Federica Gilardi, Marc Augsburger, Aurelien Thomas, Tony Fracasso, Tatjana Sajic

## Abstract

**Purpose:** Traumatic brain injury (TBI), including pediatric abusive head trauma (AHT), is the leading cause of death and disability in children and young adults worldwide. The current understanding of trauma-induced molecular changes in the brain of human subjects with intracranial haemorrhage (ICH) remains inadequate and requires further investigation to improve the outcome and management of TBI in the clinic. Calcium-mediated damage at the site of brain injury has been shown to activate several catalytic enzymes.

**Experimental design:** Serine hydrolases (SHs) are major catalytic enzymes involved in the biochemical pathways of blood coagulation, systemic inflammation and neuronal signaling. Here we investigated activity-based protein profiling (ABPP) by measuring the activity status of SH enzymes in the serum of infants with severe ICH as a consequence of AHT or atraumatic infants who died of sudden infant death syndrome (SIDS).

**Results:** Our proof-of-principle study revealed significantly reduced physiological activity of dozens of metabolic SHs in the serum of infants with severe AHT compared to the SIDS group, with some of the enzymes being related to neurodevelopment and basic brain metabolism.

**Conclusions and clinical relevance:** To our knowledge, this is the first study to investigate the ABPP of the SHs enzyme family to detect changes in their physiological activity in blood serum in severe TBI. We used antemortem (AM) serum from infants under the age of 2 years who were victims of AHT with a severe form of ICH. The analytical approach used in the proof-of-principle study shows reduced activities of serum serine lipases in AHT cases and could be further investigated in mild forms of AHT, which currently show 30% of misdiagnosed cases in clinics.

## Article Type: Rapid Communications

TBI is a major cause of death in young people under the age of 45 and remains one of the major health and socioeconomic problems worldwide[1–3]. Trauma-induced changes in the catalytic activity of enzymes involved in major biochemical cascades lead to neuroinflammation, cognitive dysfunction and increased mortality[4–8], offering potential targets for therapeutic interventions. It has been shown that calcium-mediated damage at the place of brain injury[3, 9] leads to the activation of proteolytic enzymes, as exemplified by the cysteine proteases Calpains 1/2[10] or Ubiquitin carboxy-terminal hydrolase L1 (UCHL1)[10], which latter was recently approved by the Food and Drug Administration (FDA) as a biomarker of TBI[11]. Serine proteases, which belong to the large enzyme superfamily of SHs, have been implicated as key regulators of biochemical proteolytic pathways of blood coagulation and inflammation[6, 12]. SHs of the brain have fundamental roles in central nervous system (CNS) signaling[8, 13] specifically endocannabinoid system[14, 15] via metabolic processing of major neurotransmitters and lipid messengers and thus are also important in TBI[4, 8]. Furthermore, pharmacological modulation of monoacylglycerol lipase (MAGL) activity, a key SH enzyme degrading the endocannabinoid 2-arachidonoylglycerol (2-AG) in astrocytes[14], exerts neuroprotective effects against brain trauma[4, 14, 15].

AHT is one of the most severe forms of TBI in young children, with higher mortality and morbidity than accidental injury[16–19]. Our recent study demonstrated that the SHs of complement cascades and those involved in neurodevelopment, such as Lecithin-cholesterol acyltransferase (LCAT) with their substrate lipoproteins (e.g., APOE, CLU)[20–22] known to modulate TBI[22], were highly elevated enzymes in infant victims of AHT with severe ICH compared with atraumatic cases[19].

ABPP technology uses active site-directed chemical probes to selectively target and quantify active forms of the studied enzymes [23], but not necessarily their biological amount[24]. Indeed, ABPP has high potential to capture and identify increased enzymatic activity of neuronal serine lipases involved in the metabolism of neurotransmitters[13] or those serine protease of complement and coagulation cascades[24, 25] involved in inflammation following TBI[5, 7, 26–28]. Curiously, ABPP has not yet been explored in human TBI samples for the development of diagnostic tools or eventual drug targets based on the biological activity of enzymes following TBI. Yet, the single protease activity of thrombin[6] or calpain 1 has already been explored via specific substrates or activity based sensors[10, 29], confirming that enzyme biological activity, but not protein abundance, plays a key role in molecular changes shortly after brain trauma. Currently, profiling of enzyme activity states by large-scale ABPP is usually performed in full cell lysates or in tissue homogenates[13, 23, 30] at physiological pH. Here, our goal was to evaluate the application of functional proteomics technology, ABPP, for the monitoring of enzyme activity states in the blood serum or plasma of severe TBI cases. Based on the chemical control of the clotting process, the molecular composition of blood-derived specimens differs in clinics[31]. The most frequent plasma sampling requires collector tubes that can be filled with chemical additives – anticoagulants (e.g., EDTA, heparin, citrate) – in order to prevent blood coagulation. One of the most common additives in plasma collector tubes is EDTA (i.e., ethylenediaminetetraacetic acid). EDTA chelates calcium and prevents proteolytic activity of coagulation factors involved in clotting, but can also potentially inhibit other catalytic enzymes. Another commonly used blood liquid, serum, can be collected and generated in the tubes “free of additives” via centrifugation after removal of the fibrin clot and blood cells. Hence, the choice of blood collection tubes represents an important step for each blood-based omics analysis[31, 32], including functional proteomics studies.

To measure SH activity in the bloodstream of AHT infants below the age of 24 months and compare it with thoroughly matched atraumatic controls, we apply the functional proteomics approach ABPP (**Figure 1A**) that uses a chemical probe – 6-N-biotinylaminohexylisopropylphosphorofluoridate – solution known as Fluorophosphonate (FP-1, **Figure S1A**) to selectively tag the physiologically active SHs[30, 33]. Of note, chemical specificity of this molecular probe toward serine nucleophile of enzymatically active SHs was extensively studied and described elsewhere[33]. First, we perform protocol adaptation in standard serum and plasma of atraumatic cases. We used 50 µL of control human plasma generated via tube collectors containing EDTA and the serum generated in additive-free tubes. We diluted both specimens to 1 ml with a phosphate-buffered saline (PBS) solution, proceeding with sample incubation with 5 µM/ µL of FP-1 probe that integrates biotin tag in the control positive (CP) experiment. To track non-specific bindings on the streptavidin coated beads commonly described in this type of affinity-based proteomic experiment[30], we used the control negative (CN) experiment. In control negative experiment we incubated the respective specimens with DMSO reaction solvent in which the FP-1 biotin probe was left out. All sample variants were processed in the same way (**Figure 1A**). Reassuringly, we identified almost no SH enzymes in control negative experiments of test samples, and we captured small number of nonspecific proteins in both sample types (42 and 33 proteins in CN plasma and CN serum, respectively; **Figure S1B)**. We observed that a smaller number of proteins was captured in the control positive experiment of plasma compared with the control positive experiment of serum specimen (i.e., 129 vs. 312 proteins, respectively; **Figure S1B**). As expected, the EDTA chelating agent interferes with probe binding to the catalytic enzyme, resulting in a lower number of captured SHs in the plasma compared with that in the serum (i.e., 15 vs. 54/ SHs, respectively, **Figure S1B**). Remarkably, we had captured and identified more than 50 active SH enzymes (i.e. FP-1 labeled SH) in the workflow with serum specimens (**Table S1**).

**Figure 1.**
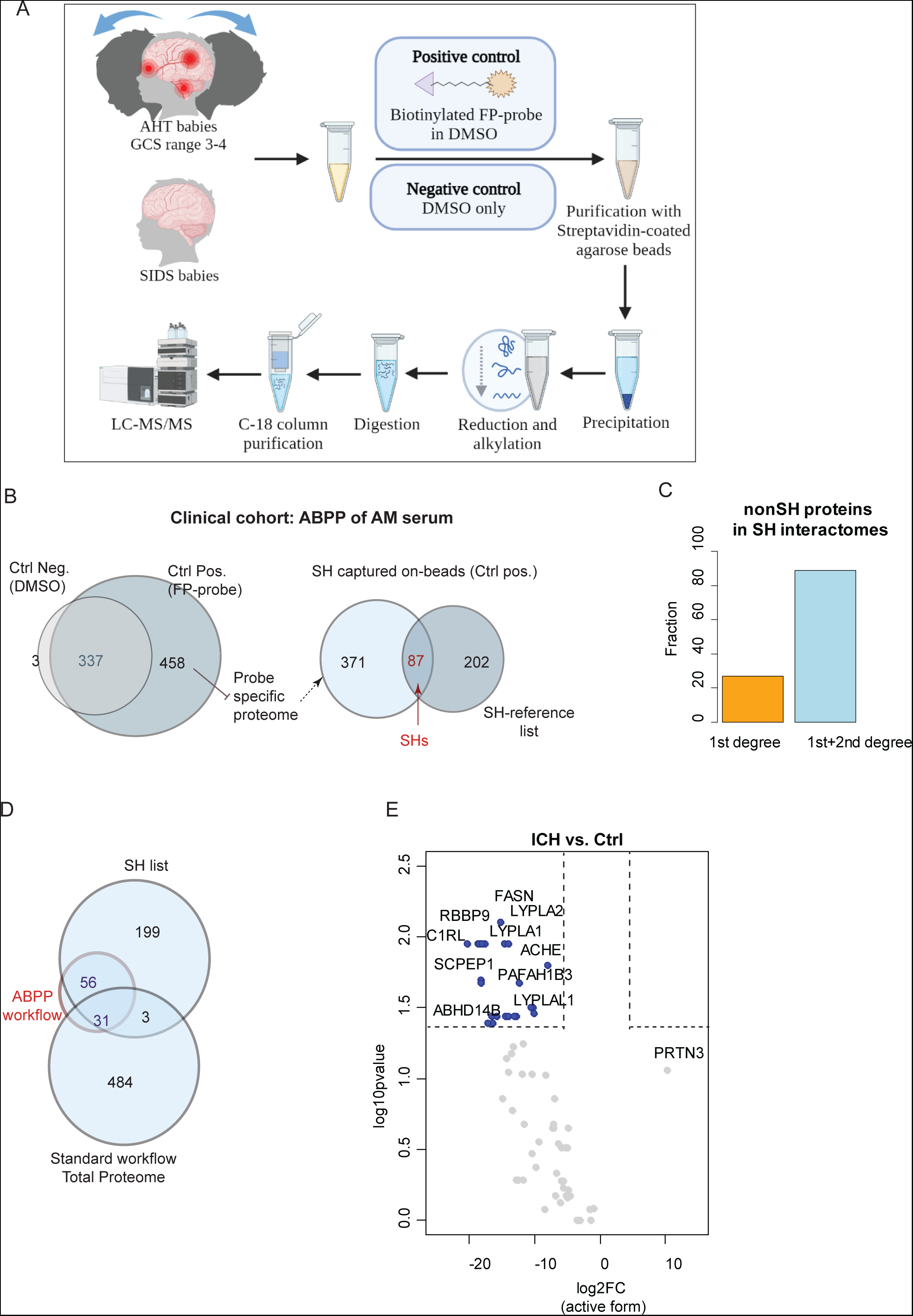
Activity based protein profiling of SHs in the serum of AHT cases and atraumatic controls. A, Blood-derived infant specimens are incubated with FP-biotin probe (control positive, CP) or reaction solvent (control negative, CN) and the active serine hydrolases are captured on streptavidin beads. Proteins captured on streptavidin beads are trypsin digested and peptides analysed by DDA-MS. B, Venn diagrams represent: an overlap of proteins captured on beads from CN and CP experiment (left) and the number of annotated SHs labelled with FP probe and captured in CP experiment (right) C, Barplots represent the fractions of proteins captured on beads annotated as first (orange) or second (blue) degree protein interactors with one of the SH enzymes. D, Venn diagram showing the overlap of SHs detected by ABPP and the standard proteomics protocol. SH list corresponds to our manually compiled list of SH enzymes from literature searches. E, Volcano plot showing comparison of FP-labelled SHs intensities log-transformed from AHT (ICH, n=5) and SIDS (sera, n=5) sera. Fold change (FC) and P-value corresponds to two-sided Wilcoxon test.

We further analysed the AM serum of ten infants via an identical ABPP workflow, five of which infants were victims of AHT with diagnosed severe ICH, while the remaining five were atraumatic controls who had died of SIDS (**Table S2**). We ensured that all the serum specimens of the respective infants were collected at Intensive Care Units (ICUs) in antemortem condition (longest delay between serum collection and death was 75 hours) and identically processed in the ABPP experiment. Remarkably, we detected 87 SHs in the AM serum of control positive experiments incubated with a chemical probe, indeed confirming affinity-based enrichment of the SH enzyme family via our workflow (**Figure 1B, Table S3**). Of note, a low number of SHs was detected sporadically in the control negative experiments, but mostly with single peptide per enzyme. For instance, the same enzymes labeled with FP1 (i.e., CP experiment) were detected with 5 to 60 peptides on average (e.g. FASN, **Figure S1C**). In addition, according to a reference integrated interaction database (IID), the resource of protein–protein interactions in humans [34], we found that 95% of non-SH proteins captured on the beads together with active SH enzymes (i.e., 354 out of 371, **Figure 1B**, right) were characterized as 1^st^ or 2^nd^ level protein interactor with SH **(Figure 1C, Table S4)**. The number of these protein SH interactors detected within “on-bead proteome digest” (i.e., 354) was higher than expected by chance (p-value < 2.2e-16, two-proportions z-test). This was confirmed through testing of 100 randomly selected protein lists of the same size (from IID database), which systematically showed a lower number of SH interactors compared with protein set enriched on beads (i.e., green arrow shows greater number of SH interactors (i.e. 354) compared to random sets <100), **Figure S1D**). For instance, we detected an overrepresentation of enzyme binding proteins and serine type endopeptidase inhibitors within proteins captured on the streptavidin beads from patient sera (Fisher Exact test FDR corrected P-value <0.01, **Figure S1E**), and this analysis indirectly confirms workflow specificity toward the SH enzyme superfamily. After comparing the results of the serum ABPP workflow with standard serum proteome profiling of the same samples for the number of enzymes detected from our manually compiled SH list (**Table S5**), we observed that the standard setup detected only 34 SHs, mainly complement proteins. ABPP workflow identified as many as 87 SH enzymes of which 56 were specifically detected via FP1-labeled SH catalytic fractions in this analytical workflow (**Figure 1D**). Furthermore, we selectively captured low-abundant serine lipases, some of which were involved in brain function and abundant in cerebrospinal fluid (CSF), lipases such as Acetylcholinesterase (ACHE)[35], Cholinesterase (BCHE)[35], PAF acetylhydrolase 29 kDa subunit (PAFAH1B3)[36] and Monoglyceride lipase (MGLL)[14] [13] (**Figure 1E**)).

Next, we focused on quantitative mass spectrometry, using confident relative MS intensities of catalytically active enzymes detectable via ABPP workflow (or LFQ intensities of FP1-labeld SH enzymes) in the serum cohort of respective infants. To our surprise, we observed that most of the detected enzymes appeared to have low physiological activity in the bloodstream of infants with severe ICH compared with the bloodstream of infants who had died of SIDS (Wilcoxon test P-value <0.05, |Log_2_FC>5|, **Figure 1E**). Only serine protease Myeloblastin (PRTN3) derived from neutrophils and involved in neutrophil-mediated matrix protein breakdown and blood-brain barrier damage[37, 38], shows a tendency for increased serum activity in severe AHT. We stress that SIDS cases had no signs of head trauma at medico-legal examination and that their specimens were processed identically to those of the AHT cases in clinical protocol as well as analytical experiments.

We then focused in the activity of serine proteases of complement cascade involved in neuroinflammation[7] and metabolic serine lipases highly expressed in brain[13]. As biochemical reactions occur at controlled stoichiometric proportions, we hypothesized that after the blood has clotted, the native serum liquid, that is free of additives, potentially contains the active proteases of complement if the latter were present in excess in the bloodstream, as we previously demonstrated in infants with intracranial bleeding induced by AHT[19]. We focused on the five complement proteases (e.g. C1S, C1R, C2, C1RL, MASP1, **Figure S2A**) that were significantly elevated in sera from severe AHT cases and for which expression levels were measured in our previous study[19]. We observed that proteolytic activities of five complement proteases did not follow their high circulating levels or correlate with markers of neuronal damage (i.e. BASP1, NCHL1, ENPP2)[19]. This is interesting because the proteases of the classical complement pathway are a key driver of secondary brain injury following intracranial haemorrhage[5, 7, 26–28].[39]Due to serum derivation via the the clotting process, which may also affect the activity of complement members, the results should be further verified in other patients with severe ICH. However, these data demonstrate that high levels of serum complement proteases in severe TBI are not necessarily representative of their biochemical function.

Remarkably, we detected that dozens of hydrolytic enzymes, mainly lipases (e.g. LYPLA2, MGLL, LCAT, **Figure 2A**), display significantly reduced hydrolytic activity in the serum of AHT infants with ICH compared with the atraumatic controls. Severe cases of AHT systematically show lower levels of active enzymes and also reduced numbers of detected peptides from FP1-labeled SHs on the streptavidin beads (**Figure 2B**). For example, metabolic SHs, such as Platelet-activating factor acetylhydrolase IB subunit alpha1 (PAFAHB13), MAGL and Acyl-protein thioesterases 1 and 2 (LYPLA1/2), displayed low catalytic activities in the serum of infants with severe ICH (**Figure 2A-B**). Furthermore, the major neuronal lipases Acetylcholinesterase (ACHE) that physiologically absorb neurotransmitter Acetylcholine[40] display significantly reduced serum activity in severe AHT cases (**Figure 2C**) compared with atraumatic controls. Interestingly, increased levels of brain-derived lipids, were previously observed in the serum of a large cohort of TBI patients via lipidomic analysis[41, 42], which we assume could also affect the lipase status in the serum of infants victims of severe AHT. Due to changes in the molecular composition of the serum, the metabolic activity of the enzymes in the serum may vary over time after injury, depending on the type and severity of the brain injury[42]. Indeed, the reduced cortical cholinergic function that we detected here in victims of severe forms of AHT through reduced ACHE activity is well known in TBI cases and monitored in serum for prognostic purposes[43].

**Figure 2.**
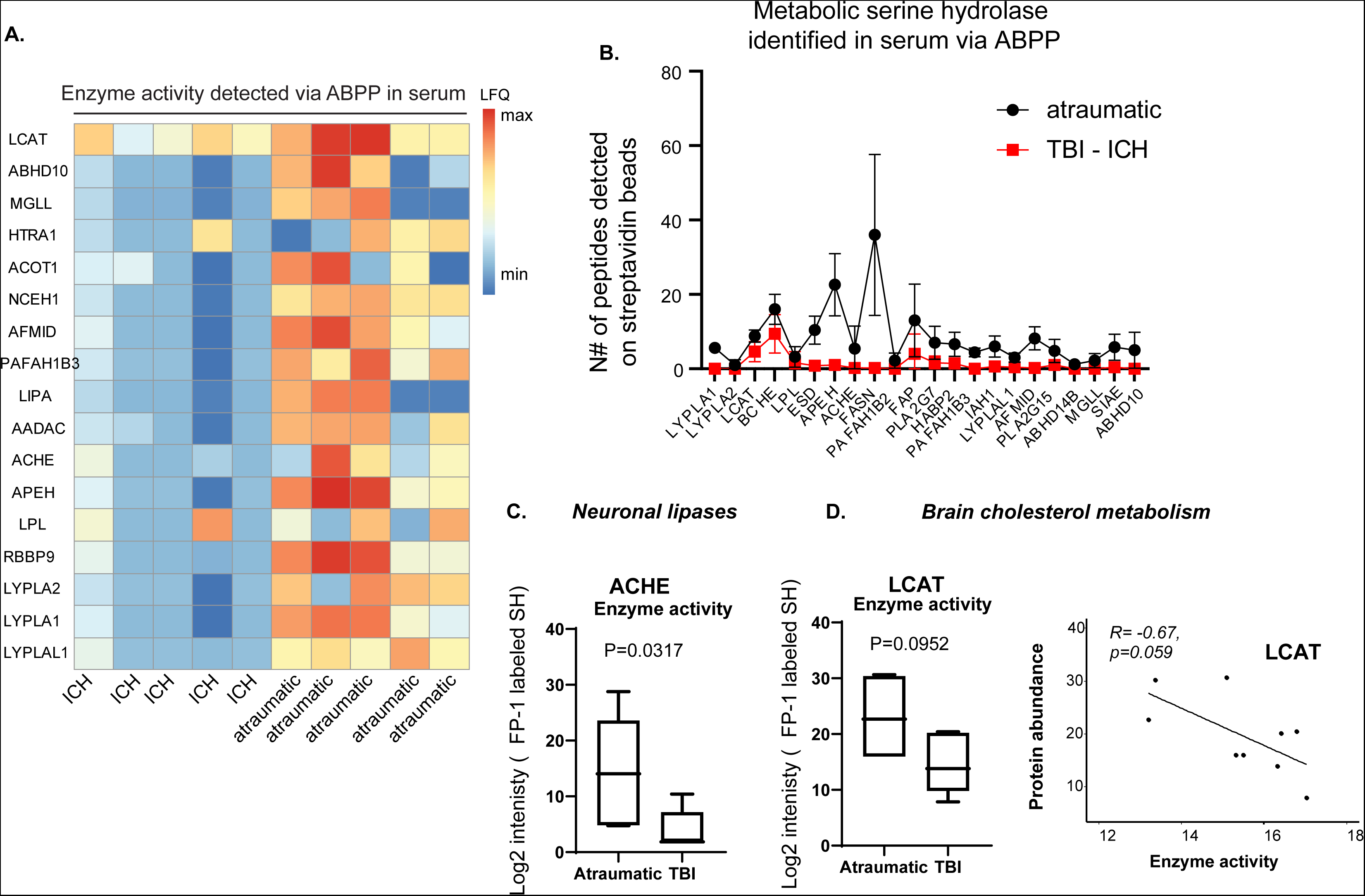
Reduced enzymatic activity of metabolic serine hydrolases in serum of AHT cases compared to SIDS. A, Heat map of FP1-labelled metabolic SHs detected in sera from ICH and atraumatic cases. FP1 probe is used for active site binding of catalytically active SH enzymes. Data correspond to scaled LFQ intensities. B, Number of peptides detected per FP1-labelled enzyme in ICH (red) and atraumatic (black) cases. C, Boxplots represent detected ACHE activity in sera of two sample groups based on LFQ intensities of FP1-labelled enzyme form. D, Boxplots represent detected LCAT enzyme activity in the corresponding sera samples. LCAT activity is estimated from the amount of FP-labelled enzyme form. P-value corresponds to two-sided Wilcoxon test. Whiskers on box plots indicate minimum to maximum observation. Spearman rho correlations of degree of LCAT catalytic activity with total serum LCAT protein level.

Along the same lines, Phosphatidylcholine-sterol acyltransferase (LCAT), in part produced in the brain by primary astrocytes[21] and highly elevated in the circulation of infant victims of AHT with severe ICH[19], also displays reduced enzymatic function. Curiously, reduced activity of LCAT enzyme in serum do not correlate with LCAT protein levels significantly elevated in severe TBI/AHT cases (**Figure 2D, Figure S2B**). LCAT esterifies free cholesterol on nascent APOE-containing lipoproteins secreted from glia and influences CSF APOE– and APOA1 levels[21]. Interestingly, several Apolipoproteins were elevated in the serum of AHT cases[19]. Glycerophospholipids are the major component of biological membranes in the brain[44], and it has been shown that an increase in CSF glycerophospholipids and lipoproteins modulates LCAT enzyme activity to maintain cholesterol homeostasis in the brain of patients with neurodegenerative diseases.[45, 46] Dysregulated lipid profiles in serum have been also associated to increased permeability of the blood brain barrier (BBB) and patient outcome after TBI[41, 42, 44]. However, the biological activity of enzymes that absorb those lipids or modulate endocannabinoid signaling following trauma[4] is poorly understood in TBI patients and needs further investigation. We assume that the status of lipases in serum could be affected by altered serum lipid composition as a result of the ability of lipids to diffuse through the BBB[42, 44] and their enzyme kinetics need to be further explored within a different time range after head injury.

Our data showed that hardly any of the enzymes were active in severe AHT/ICH cases compared to atraumatic controls (**Figure 1E**, **Figure 2A-B**). It is most likely that molecular interactions or post-traumatic biophysical changes (e.g., blood pH, temperature) that evolve over time after injury are responsible for the dysregulated enzyme activities in the serum of severe AHT cases. We expect that the metabolic activity of the enzymes that can be measured in serum using our approach will serve as good indicators of serum changes in patients with head injury or ICH, particularly in complicated forms of paediatric AHT, which are often misdiagnosed in clinical practice[16]. In contrast to severe cases of AHT (Glasgow Coma Scale score, 3–9), moderate/mild forms of AHT in infants (Glasgow Coma Scale score, 9–15) are frequently misdiagnosed because caretakers often provide inaccurate histories[17, 18, 47] and infants present symptoms non-specific to TBI[16]. Currently, no screening test exists to help clinicians in their investigations of young victims of AHT with such symptoms. The main biochemical tools in TBI management are based on standard protein measurements via the ELISA test, which could often be inaccurate. Indeed, as our results showed, serum protein levels do not always reflect the state of physiological protein function (e.g. LCAT, **Figure 2D, Figure S2B**), and this further contributes to the high level of inconsistency in the literature, especially for serum markers of paediatric TBI.[2, 48]. Other molecular hypotheses need to be explored using tools of chemical biology. Nevertheless, this technical report demonstrates that the ABPP technology needs to be further evaluated in other serum samples from TBI cases as a unique opportunity to improve clinical monitoring of complicated TBI cases by measuring serum protein function in addition to circulating expression levels.

## EXPERIMENTAL PROCEDURES

### Samples

The antemortem serum collected in the ICU from AHT cases (n=5) and atraumatic controls (infants died of sudden infant death syndrome, n=5) were collected from 2013 through 2018 in medico-legal context (**Table S2**). For ABPP experiments including protocol development, we used two different blood matrices, plasma in EDTA tube from adult human subject; and antemortem sera in free-additives tubes from adult human subject and from infants of the case-control retrospective study (Project ID 2021-01304). AHT cases were evaluated according to the guidelines of the french High Authority of Health[49]. The case-control retrospective study (Project ID 2021-01304) has been approved by the Research Ethics Committee of Geneva, Switzerland.

### ABPP protocol

Defrosted 50 µl of serum or plasma were added to 1 ml of phosphate-buffered saline (PBS) (#P-3813, Sigma). In control positive experiment (CP), we add 2 µl of 6-N-biotinylaminohexylisopropylphosphorofluoridate (FP) probe solution (Toronto Research Chemicals - Cas number 353754-93-5, TRC) to diluted serum specimens. To verify nonspecifically bind proteins to streptavidin beads we used control negative experiment (CN), and incubated the identical amount of diluted serum with 2 µl of dimethyl sulfoxide (DMSO). All samples were gently vortexed and incubated at Room Temperature (RT) for 2 hours. After incubation, 5 µl of Triton X-100 (#T8787, Sigma Aldrich) was added at 0.5% (v/v) and solutions were rotated at RT for 1 hour. Then, all solutions were desalted using PD MidiTrap G-25 (GE Healthcare) columns according to manufacturer instructions, solubilized with 75 µl (0.5% w/v) Sodium dodecyl sulfate (SDS), vortexed and incubated at 90°C for 10 minutes. After heating at 90°C, 1.5 ml of solutions were added to 70 µl of prewashed avidin beads suspension (#20347, Thermo Pierce) andwere rotated at RT for 1 hour. Then, beads were pull down at 1400 rpm at 4°C for 3 minutes, while supernatant was discarded. Beads were washed 2 times with: 1 ml SDS 1%, 1ml Urea 6M, 1 ml PBS 0.01M. For each wash, solutions were rotated at RT for 4 minutes, centrifugated at 1400 rpm for 1 minute and the supernatant was discarded. The on-bead protein digestion was performed in 6M UREA buffer. We add 7 µl tris(2-carboxyethyl)phosphine (TCEP) (Sigma) to streotavidin coated beads and incubated samples at RT at 60 rpm for 30 minutes. Next, 12 µl iodoacetic acid (IAA) (Sigma) was added to the solutions and samples were incubated at RT at 60 rpm for 30 minutes. Solutions were centrifugated for 3 minutes at 1400 rpm and the supernatant was discarded. Beads were washed 3 times with 500 µl Urea 1M, and each time centrifugation was performed for 1 minute at 1400 rpm. Finally, we add 6 µl of Trypsin (Promega) to each sample for protein digestion overnight at 100 rpm at 37°C in 400 µl 1M Urea. The next day, digestion was stopped with 10 µl of 10% formic acid, the solutions were vortexed and centrifuged at maximum speed for 10 minutes to collect supernatants containing digested peptides. After peptide clean-up on C-18 columns (Nest Group Inc SEM SS18V MICROSPIN COLUMNS C18 SILICA), the cleaned peptides were eluted with 200 µl of buffer B (1% formic acid, 5% H2O in ACN), centrifuged at 2000 rpm for 2 minutes, and the excess buffer B was evaporated using a SpeedVac (CentriVap® concentrator). Pellets were resuspended with 25 µl Buffer A (1% Formic Acid, 5% Acetonitrile (ACN) in H2O), vortexed at maximum speed for 30 minutes with incubation at 37°C. Before performing LC-MS analysis, samples that underwent ABPP-protocol were centrifugated at maximum speed for 15 minutes and used for peptide quantification using a NanoDrop UV Spectrophotometer (ThermoFisher Scientific) and absorbance was measured at 280 nm.

### Liquid chromatography-electrospray ionization tandem mass spectrometry (LC-ESI-MS/MS)

Samples of collected peptides digests were vortexed and centrifuged at maximum speed for 10 minutes. 1 µl of retention time (rt) peptides (iRT-kit, Biognosys) were added to the solutions. Solutions were analyzed with LC-ESI-MS/MS workflow on an Orbitrap Fusion Lumos Tribrid mass spectrometer (Thermo Fisher Scientific) equipped with an Easy nLC1200 liquid chromatography system (Thermo Fisher Scientific). Peptides were trapped on an Acclaim pepmap100, C18, 3μm, 75μm x 20mm nano trap-column (Thermo Fisher Scientific) and separated on a 75 μm x 500 mm, C18 ReproSil-Pur (Dr. Maisch GmBH), 1.9 μm, 100 Å, home-made column. The analytical separation was run for 90 min using a gradient of H2O/FA 99.9%/0.1% (solvent A) and CH3CN/FA 80%/0.1% (solvent B). The gradient was run as follows: 0-5 min 95 % A and 5 % B, then to 65 % A and 35 % B in 60 min, and finally to 5% A and 95% B in 10 min with a stay for 15 min at this composition. Flow rate was of 250 nL/min. Data-dependant analysis (DDA) was performed with MS1 full scan at a resolution of 120’000 FWHM followed by as many subsequent MS2 scans on selected precursors as possible within 3 second maximum cycle time. MS1 was performed in the Orbitrap with an AGC target of 4 x 105, a maximum injection time of 50 ms and a scan range from 400 to 2000 m/z. MS2 was performed in the ion-trap with an AGC target at 1 x 104 and a maximum injection time of 35 ms. Isolation windows was set at 1.2 m/z and 30% normalised collision energy was used for higher-energy collisional dissociation (HCD). Dynamic exclusion was set to 20s.

### Analysis of generated MS data

MS raw data files were analyzed with MaxQuant proteomic software (version 2.1.3.0)[50]. For peptide search and generation of quantitative data matrices via label free quantification (LFQ), default parameters included trypsin digestion were used. In brief our parameters allowed two missed cleavage events, ‘Carbamidomethyl (C)’ was chosen as fixed modifications and ‘Oxidation (M)’ as variable modifications. We selected option “Re-quantify” for first search as calibration steps and “Match between runs” for association of spectral identification across LC-MS/MS runs based on RT and accurate peptide mass. The mass tolerances were set to 20 ppm for precursor-ions and 0.1 Da for fragment-ions. Unique proteins identified at 1% of protein false discovery rate (FDR) were used to generate the list of proteins from each individual sample. Human Uniprot IDs were used as template (2022.10.18-04.39.06.52). Immunoglobulins and contaminants were excluded. We manually compiled a list of 289 SH enzymes based on literature searches, using, for example, the Merops database (https://www.ebi.ac.uk/merops/) for serine protease annotation, and a previously published list of metabolic serine hydrolases.[13, 51]. For the generation of protein data matrices we used Maxquant output data file were imported in R software (version 2022.02.3) and data were processed using MSstats package[52]. The function “dataProcess” was used and imputed arguments were the following: log2-transformation of the data, quantile normalized at peptide levels, top3 peptide features with 2 minimal feature count per protein, decomposition of the original matrix using Tukey’s median polish (TMP) and missing values were censored at 0.999 maximum Quantile. After data processing, proteins were quantified using “quantification” function and batch effect was estimated using non-scaled clustering with Euclidean distances with pheatmap R-package. Batches were systematically matched to the time of samples injection in LC-MS/MS experiments. Batch effect was corrected at peptide level using ComBat as discrete and Loess regression as continuous function using proBatch package[53]. Batch effect correction was performed on abundance intensities of peptides after MSstat normalization of data which were not censored and not imputed. Abs threshold was implemented at 5 and pct threshold at 0.20. After batch effect correction, proteins were reconstituted from peptides with “dcast” function from reshape2 package with average of the corresponding peptide.

### Statistical Analysis and data visualization

Log2 transformed protein data matrix after batch effect correction was used for statistical analysis. The Wilcoxon test was used to compare the level of enzyme activity between groups and P value < 0.05 was considered significant. For correlation analyses, Spearman correlation was calculated. The fraction of proteins detected on the CP beads (i.e. 371 proteins detected on the FP-probe-beads conjugate) was analysed via the IID database[34] for binary interactions with SH enzymes. Of these, 354 proteins were annotated as either 1^st^ or 2^nd^ degree SH interactors.

To exclude experimental bias (due to the large number of SH interactors detected on beads), we also check the frequency of SH captured protein interactors found in other 100 random interactomes (i.e. 100 protein lists generated by randomly selecting protein interactors from IID). The number of physical interactions for 100 random protein subsets generated by sampling random proteins was visulaised by histogram plot. Green arrow indicates number of confirmed physical interactions found within subsets of 354 proteins captured on beads. GO analysis of molecular function (Fisher exact test) was performed on 106 proteins detected on positive control beads and annotated as 1st degree protein interaction with SH enzymes. The most important GO molecular annotations were classified according to the FDR corrected P-value significance level (FDR<0.001) and the magnitude of fold enrichment from overrepresentation test. Venn diagrams, generated using Venny software (Venny 2.1.0 - BioinfoGP), were used to verify the overlap of proteins detected by different protocols.

## Acknowledgments

We are grateful to Alexandre Hainard and all members at the Proteomics Core Facility at the Faculty of Medicine at the University of Geneva for their excellent support.

## Data availability statement

Raw data files related to standard protein abundances generated by standard proteomics are available via ProteomeXchange with identifier PXD028801. Raw data files generated using the ABPP protocol are being submited to ProteomeXchange.

## Abbreviations

ABPP: Activity based proteome profiling
FP: Fluorophosphonate
(SH(s): Serine Hydrolase(s)
AHT: Abusive Head Trauma
ICH: Intracranial Hemorrhage
SIDS: sudden infant death syndrome

## Supporting information

### Supplementary Figures

**Supplementary Figure S1.**
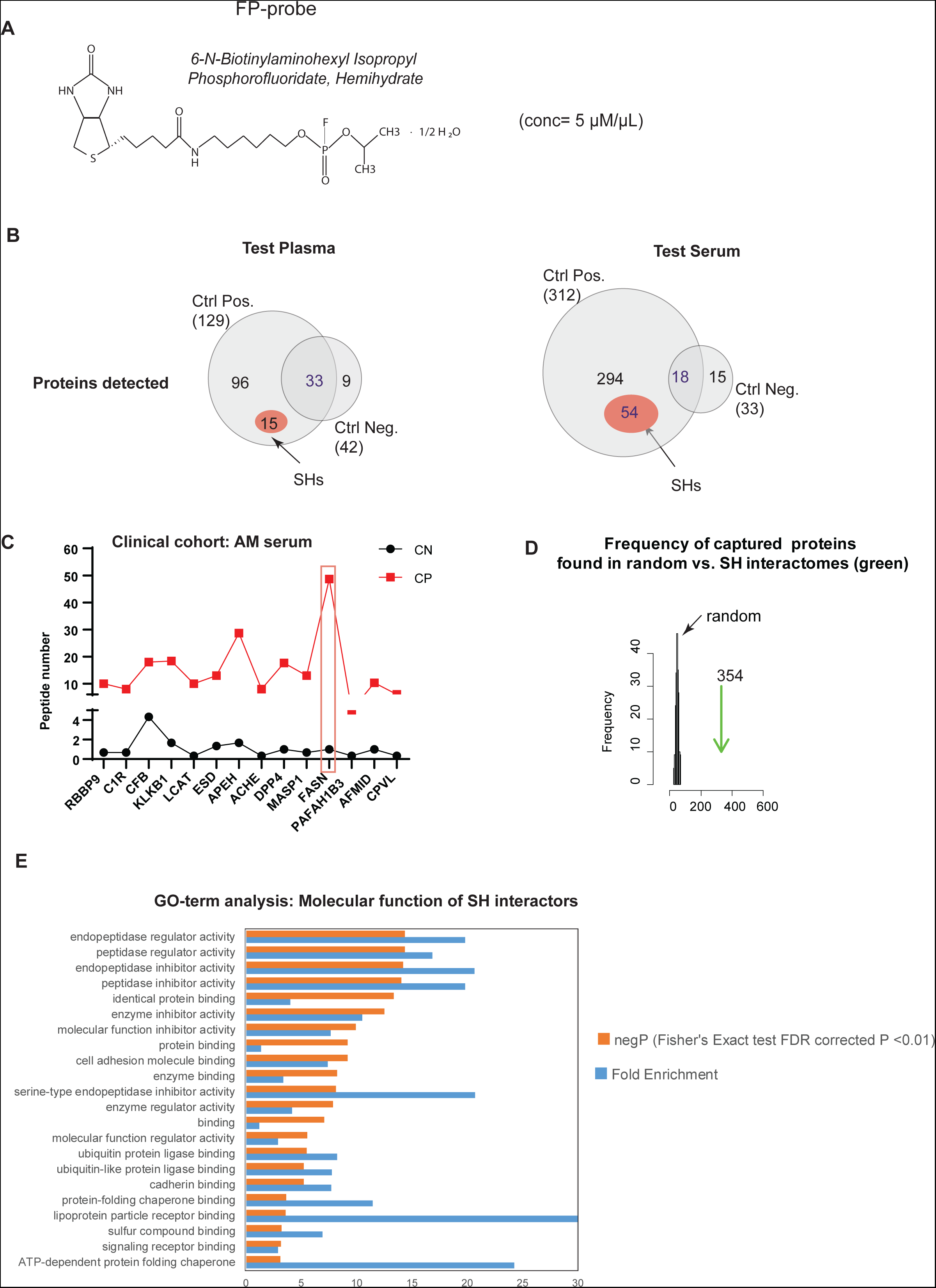
A, Structure of the fluorophosphonate (FP) biotin probe. B, ABPP protocol assay in two different human blood samples: plasma (left) and serum (right). Representative Venn diagrams show the overlap of proteins captured on beads from the CN and CP experiments (left) for each sample type, plasma and serum, respectively. The number of annotated SHs, labelled with the FP probe and captured in the CP experiment, within the total number of captured proteins of each respective sample type is shown in red circles. C, Number of peptides detected in the CP experiment per FP1-labelled enzyme (red) and per corresponding enzymes (not FP1-labelled) in the CN experiment (black). D, Frequency of proteins captured on the beads found in random interactomes compared to the number of proteins captured on the beads found in the SH interactom (green arrow). E, Most important GO molecular function detected within 1st degree SH interactors by overrepresentation in Ficher exact test with FDR corrected P-value <0.001.

**Supplementary Figure S2.**
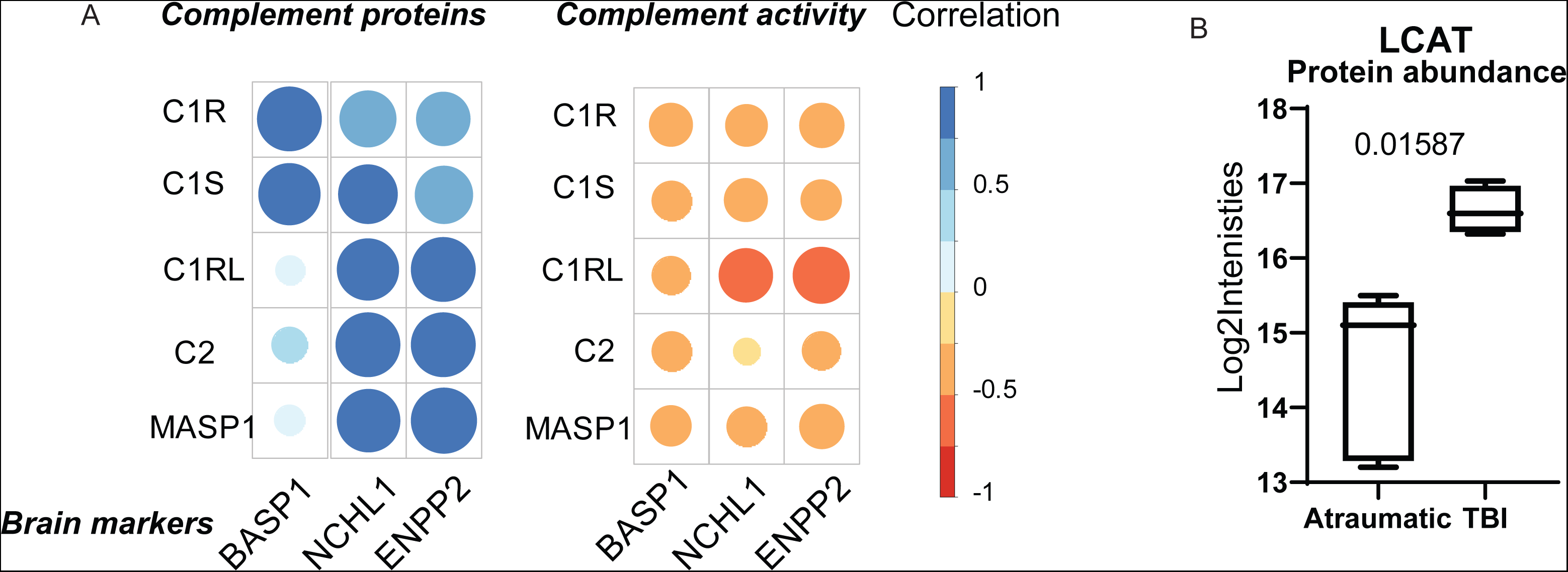
A, Spearman rho correlation analysis of serum levels of: a) five complement proteins elevated in AHT cases with three markers of brain damage (left) and b) proteolytic activity of the corresponding five complement proteins with three markers of brain damage (right). The size and colour of the circles represent either the magnitude of the correlation or the direction of the correlation (positive or negative). B, Box plots represent the detected LCAT protein abundance in the corresponding sera samples. P-value corresponds to the Wilcoxon test. Whiskers on boxplots indicate minimum to maximum observation. LCAT protein abundance available via standard proteomic digestion of corresponding infant sera.

## References

[1] Maas, A. I., Stocchetti, N., Bullock, R., Moderate and severe traumatic brain injury in adults. Lancet Neurol 2008, 7, 728–741.

[2] Marzano, L. A. S., Batista, J. P. T., de Abreu Arruda, M., de Freitas Cardoso, M. G., et al., Traumatic brain injury biomarkers in pediatric patients: a systematic review. Neurosurg Rev 2022, 45, 167–197.

[3] Sulhan, S., Lyon, K. A., Shapiro, L. A., Huang, J. H., Neuroinflammation and blood-brain barrier disruption following traumatic brain injury: Pathophysiology and potential therapeutic targets. J Neurosci Res 2020, 98, 19–28.

[4] Chen, C., Endocannabinoid control of neuroinflammation in traumatic brain injury by monoacylglycerol lipase in astrocytes. Neural Regen Res 2023, 18, 1023–1024.

[5] De Blasio, D., Fumagalli, S., Orsini, F., Neglia, L., et al., Human brain trauma severity is associated with lectin complement pathway activation. J Cereb Blood Flow Metab 2019, 39, 794–807.

[6] Gong, Y., Xi, G., Hu, H., Gu, Y., et al., Increase in brain thrombin activity after experimental intracerebral hemorrhage. Acta Neurochir Suppl 2008, 105, 47–50.

[7] Hammad, A., Westacott, L., Zaben, M., The role of the complement system in traumatic brain injury: a review. J Neuroinflammation 2018, 15, 24.

[8] Herr, D. R., Chew, W. S., Satish, R. L., Ong, W. Y., Pleotropic Roles of Autotaxin in the Nervous System Present Opportunities for the Development of Novel Therapeutics for Neurological Diseases. Mol Neurobiol 2020, 57, 372–392.

[9] Ng, S. Y., Lee, A. Y. W., Traumatic Brain Injuries: Pathophysiology and Potential Therapeutic Targets. Front Cell Neurosci 2019, 13, 528.

[10] Kudryashev, J. A., Waggoner, L. E., Leng, H. T., Mininni, N. H., Kwon, E. J., An Activity-Based Nanosensor for Traumatic Brain Injury. ACS Sens 2020, 5, 686–692.

[11] Papa, L., Brophy, G. M., Alvarez, W., Hirschl, R., et al., Sex differences in time course and diagnostic accuracy of GFAP and UCH-L1 in trauma patients with mild traumatic brain injury. Sci Rep 2023, 13, 11833.

[12] Moore, E. E., Moore, H. B., Kornblith, L. Z., Neal, M. D., et al., Trauma-induced coagulopathy. Nat Rev Dis Primers 2021, 7, 30.

[13] Viader, A., Ogasawara, D., Joslyn, C. M., Sanchez-Alavez, M., et al., A chemical proteomic atlas of brain serine hydrolases identifies cell type-specific pathways regulating neuroinflammation. Elife 2016, 5, e12345.

[14] Savinainen, J. R., Saario, S. M., Laitinen, J. T., The serine hydrolases MAGL, ABHD6 and ABHD12 as guardians of 2-arachidonoylglycerol signalling through cannabinoid receptors. Acta Physiol (Oxf) 2012, 204, 267–276.

[15] Hu, M., Zhu, D., Zhang, J., Gao, F., et al., Enhancing endocannabinoid signalling in astrocytes promotes recovery from traumatic brain injury. Brain 2022, 145, 179–193.

[16] Jenny, C., Hymel, K. P., Ritzen, A., Reinert, S. E., Hay, T. C., Analysis of missed cases of abusive head trauma. JAMA 1999, 281, 621–626.

[17] Letson, M. M., Cooper, J. N., Deans, K. J., Scribano, P. V., et al., Prior opportunities to identify abuse in children with abusive head trauma. Child Abuse Negl 2016, 60, 36–45.

[18] Narang, S. K., Fingarson, A., Lukefahr, J., Council On Child, A., Neglect, Abusive Head Trauma in Infants and Children. Pediatrics 2020, 145.

[19] Wiskott, K., Gilardi, F., Hainard, A., Sanchez, J. C., et al., Blood proteome of acute intracranial hemorrhage in infant victims of abusive head trauma. Proteomics 2023, 23, e2200078.

[20] Zhao, Y., Thorngate, F. E., Weisgraber, K. H., Williams, D. L., Parks, J. S., Apolipoprotein E is the major physiological activator of lecithin-cholesterol acyltransferase (LCAT) on apolipoprotein B lipoproteins. Biochemistry 2005, 44, 1013–1025.

[21] Hirsch-Reinshagen, V., Donkin, J., Stukas, S., Chan, J., et al., LCAT synthesized by primary astrocytes esterifies cholesterol on glia-derived lipoproteins. J Lipid Res 2009, 50, 885–893.

[22] Raulin, A. C., Martens, Y. A., Bu, G., Lipoproteins in the Central Nervous System: From Biology to Pathobiology. Annu Rev Biochem 2022, 91, 731–759.

[23] Cravatt, B. F., Wright, A. T., Kozarich, J. W., Activity-based protein profiling: from enzyme chemistry to proteomic chemistry. Annu Rev Biochem 2008, 77, 383–414.

[24] Fonovic, M., Bogyo, M., Activity-based probes as a tool for functional proteomic analysis of proteases. Expert Rev Proteomics 2008, 5, 721–730.

[25] Bachovchin, D. A., Cravatt, B. F., The pharmacological landscape and therapeutic potential of serine hydrolases. Nat Rev Drug Discov 2012, 11, 52–68.

[26] Amara, U., Rittirsch, D., Flierl, M., Bruckner, U., et al., Interaction between the coagulation and complement system. Adv Exp Med Biol 2008, 632, 71–79.

[27] Morgan, B. P., Harris, C. L., Complement, a target for therapy in inflammatory and degenerative diseases. Nat Rev Drug Discov 2015, 14, 857–877.

[28] Pedragosa, J., Mercurio, D., Oggioni, M., Marquez-Kisinousky, L., et al., Mannose-binding lectin promotes blood-brain barrier breakdown and exacerbates axonal damage after traumatic brain injury in mice. Exp Neurol 2021, 346, 113865.

[29] Kudryashev, J. A., Madias, M. I., Kandell, R. M., Lin, Q. X., Kwon, E. J., An Activity-Based Nanosensor for Minimally-Invasive Measurement of Protease Activity in Traumatic Brain Injury. Adv Funct Mater 2023, 33.

[30] Jessani, N., Niessen, S., Wei, B. Q., Nicolau, M., et al., A streamlined platform for high-content functional proteomics of primary human specimens. Nat Methods 2005, 2, 691–697.

[31] Vignoli, A., Tenori, L., Morsiani, C., Turano, P., et al., Serum or Plasma (and Which Plasma), That Is the Question. J Proteome Res 2022, 21, 1061–1072.

[32] Geyer, P. E., Voytik, E., Treit, P. V., Doll, S., et al., Plasma Proteome Profiling to detect and avoid sample-related biases in biomarker studies. EMBO Mol Med 2019, 11, e10427.

[33] Liu, Y., Patricelli, M. P., Cravatt, B. F., Activity-based protein profiling: the serine hydrolases. Proc Natl Acad Sci U S A 1999, 96, 14694–14699.

[34] Kotlyar, M., Pastrello, C., Malik, Z., Jurisica, I., IID 2018 update: context-specific physical protein-protein interactions in human, model organisms and domesticated species. Nucleic Acids Res 2019, 47, D581–D589.

[35] Giacobini, E., Cholinesterases: new roles in brain function and in Alzheimer’s disease. Neurochem Res 2003, 28, 515–522.

[36] Adachi, H., Tsujimoto, M., Hattori, M., Arai, H., Inoue, K., cDNA cloning of human cytosolic platelet-activating factor acetylhydrolase gamma-subunit and its mRNA expression in human tissues. Biochem Biophys Res Commun 1995, 214, 180–187.

[37] Aube, B., Levesque, S. A., Pare, A., Chamma, E., et al., Neutrophils mediate blood-spinal cord barrier disruption in demyelinating neuroinflammatory diseases. J Immunol 2014, 193, 2438–2454.

[38] Santos-Lima, B., Pietronigro, E. C., Terrabuio, E., Zenaro, E., Constantin, G., The role of neutrophils in the dysfunction of central nervous system barriers. Front Aging Neurosci 2022, 14, 965169.

[39] Perez-Riverol, Y., Bai, J., Bandla, C., Garcia-Seisdedos, D., et al., The PRIDE database resources in 2022: a hub for mass spectrometry-based proteomics evidences. Nucleic Acids Res 2022, 50, D543–D552.

[40] Silman, I., Sussman, J. L., Acetylcholinesterase: how is structure related to function? Chem Biol Interact 2008, 175, 3–10.

[41] Thomas, I., Dickens, A. M., Posti, J. P., Czeiter, E., et al., Serum metabolome associated with severity of acute traumatic brain injury. Nat Commun 2022, 13, 2545.

[42] Oresic, M., Posti, J. P., Kamstrup-Nielsen, M. H., Takala, R. S. K., et al., Human Serum Metabolites Associate With Severity and Patient Outcomes in Traumatic Brain Injury. EBioMedicine 2016, 12, 118–126.

[43] Zhang, Q. H., Li, A. M., He, S. L., Yao, X. D., et al., Serum Total Cholinesterase Activity on Admission Is Associated with Disease Severity and Outcome in Patients with Traumatic Brain Injury. PLoS One 2015, 10, e0129082.

[44] Nessel, I., Michael-Titus, A. T., Lipid profiling of brain tissue and blood after traumatic brain injury: A review of human and experimental studies. Semin Cell Dev Biol 2021, 112, 145–156.

[45] Demeester, N., Castro, G., Desrumaux, C., De Geitere, C., et al., Characterization and functional studies of lipoproteins, lipid transfer proteins, and lecithin:cholesterol acyltransferase in CSF of normal individuals and patients with Alzheimer’s disease. J Lipid Res 2000, 41, 963–974.

[46] Vitali, C., Wellington, C. L., Calabresi, L., HDL and cholesterol handling in the brain. Cardiovasc Res 2014, 103, 405–413.

[47] Colombari, M., Troakes, C., Turrina, S., Tagliaro, F., et al., Spinal cord injury as an indicator of abuse in forensic assessment of abusive head trauma (AHT). Int J Legal Med 2021, 135, 1481–1498.

[48] Glushakova, O. Y., Glushakov, A. V., Hayes, R. L., Finding effective biomarkers for pediatric traumatic brain injury. Brain Circ 2016, 2, 129–132.

[49] Shaken Baby Syndrom or non-accidental head injury caused by shaking : Clinical Practice Guidelines, French High Authority of Health. 2017.

[50] Cox, J., Mann, M., MaxQuant enables high peptide identification rates, individualized p.p.b.-range mass accuracies and proteome-wide protein quantification. Nat Biotechnol 2008, 26, 1367–1372.

[51] Long, J. Z., Cravatt, B. F., The metabolic serine hydrolases and their functions in mammalian physiology and disease. Chem Rev 2011, 111, 6022–6063.

[52] Choi, M., Chang, C. Y., Clough, T., Broudy, D., et al., MSstats: an R package for statistical analysis of quantitative mass spectrometry-based proteomic experiments. Bioinformatics 2014, 30, 2524–2526.

[53] Cuklina, J., Lee, C. H., Williams, E. G., Sajic, T., et al., Diagnostics and correction of batch effects in large-scale proteomic studies: a tutorial. Mol Syst Biol 2021, 17, e10240.

